# Quantitative mapping of dense microtubule arrays in mammalian neurons

**DOI:** 10.1101/2021.02.26.432992

**Authors:** Eugene A. Katrukha, Daphne Jurriens, Desiree Salas Pastene, Lukas C. Kapitein

**Affiliations:** Cell Biology, Neurobiology and Biophysics, Department of Biology, Faculty of Science, Utrecht University, Padualaan 8, 3584 CH Utrecht, the Netherlands

**Keywords:** cytoskeleton, microtubules, hippocampal neurons, STED, expansion microscopy

## Abstract

The neuronal microtubule cytoskeleton underlies the polarization and proper functioning of neurons, amongst others by providing tracks for motor proteins that drive intracellular transport. Different subsets of neuronal microtubules, varying in composition, stability and motor preference, are known to exist, but the high density of microtubules has so far precluded mapping their relative abundance and three-dimensional organization. Here we use different super-resolution techniques (STED, Expansion Microscopy) to explore the nanoscale organization of the neuronal microtubule network. This revealed that in dendrites stable, acetylated microtubules are enriched in the core of the dendritic shaft, while dynamic, tyrosinated microtubules enrich near the plasma membrane, thus forming a shell around the stable microtubules. Moreover, using a novel analysis pipeline we quantified the absolute number of acetylated and tyrosinated microtubules within dendrites and found that they account for 65-75% and ∼20-30% of all microtubules, respectively, leaving only few microtubules that do not fall in either category. Because these different microtubule subtypes facilitate different motor proteins, these novel insights help to understand the spatial regulation of intracellular transport.

## INTRODUCTION

The extended and polarized morphology of neurons is established and maintained by the cytoskeleton [1, 2]. One of the functions of the microtubule cytoskeleton is to provide a transport network inside the neuron’s long axon and branched dendrites [3, 4]. Directional transport is enabled by the structural polarity of microtubules, which is recognized by motor proteins that drive transport to either the minus end (dynein) or plus end (most kinesins). Using this network, intracellular cargos attached to microtubule-based motor proteins (kinesins and dynein) can be shipped between different neuronal compartments [5, 6]. To facilitate proper delivery, transport is regulated at multiple levels. For example, molecular motors, motor adaptor proteins and cargos themselves undergo tight biochemical regulation in response to changes in metabolic state and extracellular cues [5, 7]. An equally important component is the composition and spatial distribution of microtubule tracks, which is the main subject of this study.

Previous work has revealed that the affinity of motors for the microtubule lattice can be modulated by microtubule-associated proteins (MAPs) or by post-translational modifications (PTM) of tubulin [8-13]. For example, kinesin-1 motors move preferentially on microtubules marked by acetylation and detyrosination [14, 15], while kinesin-3 prefers tyrosinated microtubules [16-18]. When tubulin is incorporated into the microtubule lattice it carries a C-terminal tyrosine, which can subsequently be proteolytically removed to yield detyrosinated microtubules[3, 13]. Therefore, tyrosinated tubulin can be regarded as a marker for freshly polymerized microtubules. Such microtubules undergo cycles of growth and shrinkage and are therefore referred to as dynamic microtubules. Following polymerization, tubulins can also acquire new chemical groups through post-translational modifications, such as acetylation and polyglutamylation. Such modifications often accumulate on microtubules that are long-lived and resist cold-induced or drug-induced depolymerization, which are therefore termed stable microtubules.

Despite many biochemical and physiological studies underpinning the importance of various microtubule modifications [9, 13, 19, 20], little is known about the relative abundance and spatial organization of different microtubule subsets within neurons. In earlier work, we revealed that stable and dynamic microtubules in dendrites are organized differently and often have opposite orientations, explaining why kinesin-3 can drive efficient anterograde transport in dendrites, unlike kinesin-1 [17]. Nonetheless, many important aspects of the neuronal microtubule array have remained unexplored. First, do tyrosination and acetylation mark two clearly defined subsets or are there also subsets that are both highly tyrosinated and acetylated? Furthermore, what is the three-dimensional organization of different subsets and their relative abundance? Finally, do acetylation and tyrosination together mark all microtubules or are there additional subsets that carry neither of these groups? Although microtubule organization in dendrites has previously been studied using electron microscopy [21, 22], this method is difficult to combine with selective markers and therefore cannot robustly identify and map microtubule subsets throughout dendrites.

Here we use a variety of super-resolution techniques to explore the quantitative and spatial distribution of microtubule subsets in dendrites. We find that acetylated microtubules accumulate in the core of the dendritic shaft, surrounded by a shell of tyrosinated microtubules. High-resolution microscopy enabled frequent detection of individual microtubule segments, which could be used to carefully quantify the tyrosination and acetylation levels of these segments. This revealed that these two modifications are anti-correlated and define two distinct microtubule subsets. In addition, it enabled us to estimate the absolute number of acetylated and tyrosinated microtubules within dendrites, which revealed that they account for 65-75% and ∼20-30% of all microtubules, respectively, leaving only few microtubules that do not fall in either category. Together, these results provide new quantitative insights into the uniquely organized dendritic microtubule network and help to understand the spatial regulation of neuronal transport.

## RESULTS

We started by mapping the spatial distribution of acetylated and tyrosinated microtubules throughout the dendrite using both 2D and 3D stimulated emission depletion (STED) microscopy. Consistent with our earlier observations, this revealed that acetylated microtubules in DIV9 neurons tend to be distributed closer to the central axis of the dendrite, while the tyrosinated microtubules seems to be enriched at the outer surface, close to the membrane (Fig. 1A,B). To quantify this observation, we built radial distribution maps of the intensity of acetylated and tyrosinated microtubules along the dendrite, which could be averaged to quantitatively describe the radial distribution of the two subsets (Fig.1C,D, Supplementary Movie 1). This revealed that the differential spatial organization was maintained along the length of an individual dendrite (Fig.1D,E). Furthermore, we observed it in dendrites with various diameters (Fig.S1A,B), independent of STED imaging modality (Fig.1F,G).

**Figure 1.**
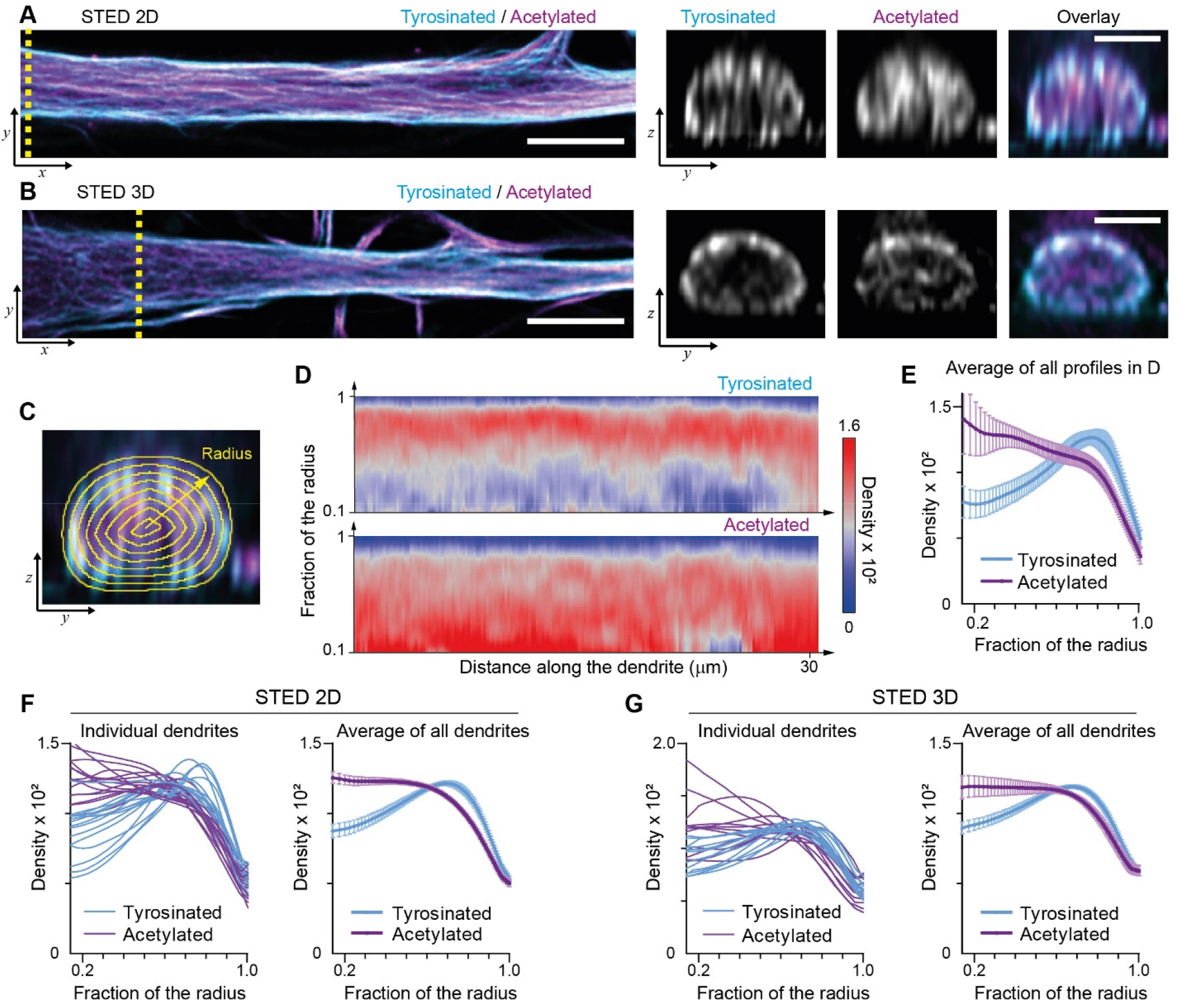
Radial distribution of microtubule subtypes in dendrites imaged using STED microscopy. (A)-(B) Representative single planes in XY (left) and YZ cross sections along the yellow dashed line (right) of a dendrite imaged with 2D (A) and 3D (B) STED. Scale bar 5 µm (XY) and 2 µm (XY). (C)Quantification of the radial intensity distribution in YZ cross sections. The outer yellow contour marks the outline of a dendrite and concentric smaller circles represent contours of smaller radius used for quantification. (D)Heatmaps of normalized radial intensity distributions for tyrosinated (top) and acetylated (bottom) microtubules along the length of the dendrite shown in (A). (E) Radial distribution of tyrosinated (cyan) and acetylated (magenta) microtubule posttranslational modifications averaged over the length of the dendrite shown in (A) (n=176 profiles). Error bars represent SD. (F)-(G) Radial distribution of modifications averaged per dendrite (left) and over all dendrites (right) imaged using 2D (panel (F), n=4971 profiles, 15 cells, N=2 independent experiments) or 3D STED (panel (G), n=5891 profiles, 12 cells, N=2 independent experiments). Error bars represent S.E.M.

We next attempted to quantify the number of microtubules for each subset. This cannot be achieved by just comparing fluorescent intensities, because staining efficiencies and fluorophore properties differ for each subset and need to be rescaled using single microtubules of each type as a reference. However, we were not able to distinguish individual microtubules within axons or dendrites, since (as it is known from electron microscopy studies) the distance between adjacent microtubules is often smaller than the resolution of STED. Microtubules in the cell body, however, were more dispersed and here individual microtubules could often be resolved (Fig 2A,B). We therefore set out to develop a work flow to enable the robust quantification of acetylation and tyrosination levels on individual microtubules, which could subsequently be used to determine the number of acetylated and tyrosinated microtubules in dendrites.

**Figure 2.**
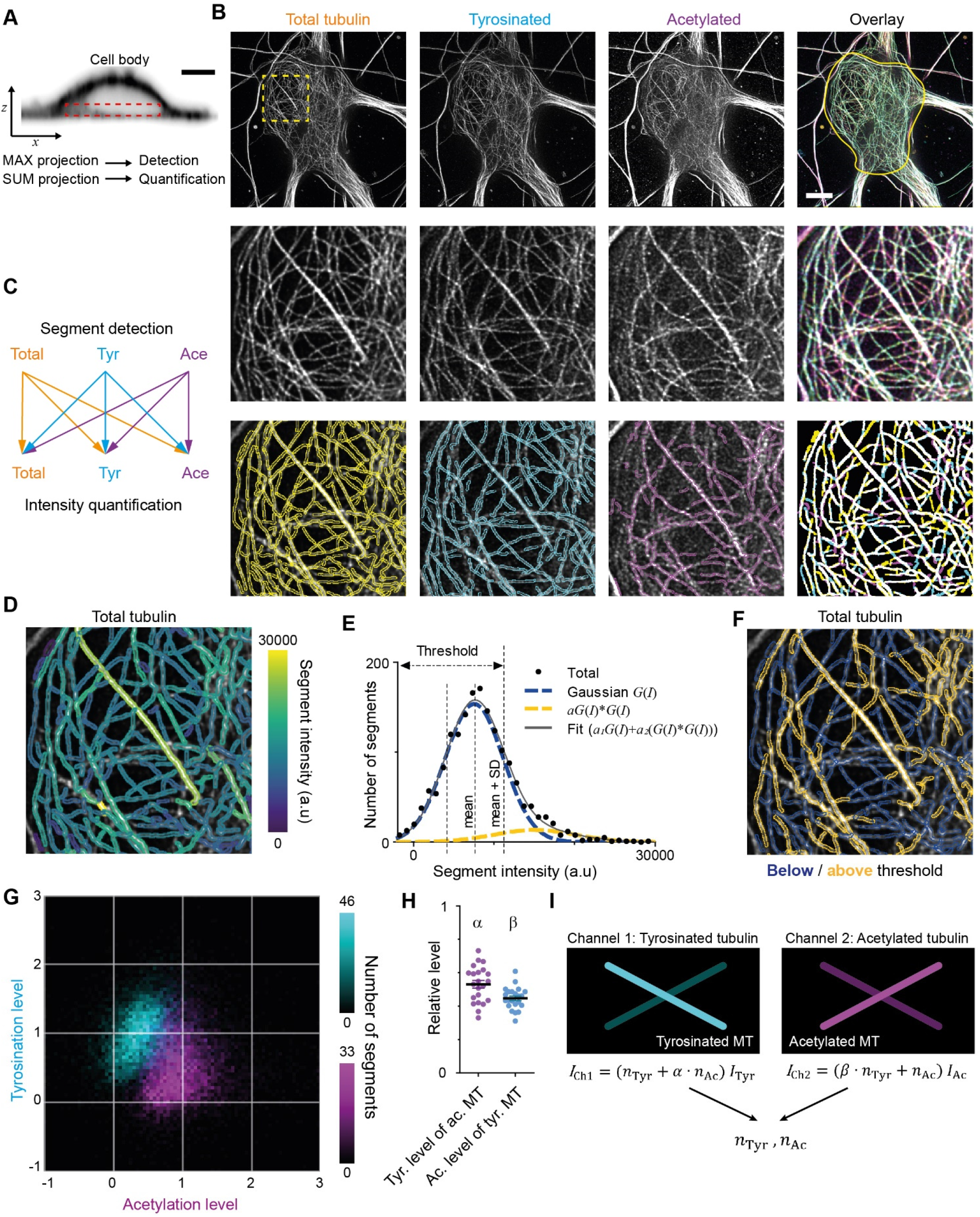
Analysis pipeline for detection and intensity quantification of individual (post-translationally modified) microtubules. (A) Vertical cross section along a neuronal cell body (soma). Dashed rectangle marks the volume (sub-z-stack) under the nucleus used for microtubule filament detection (maximum intensity projection) and quantification (sum of all slices). Scale bar 5 µm. (B) (top row) Maximum intensity projection of 2D STED z-stacks of DIV9 neurons stained for alpha-tubulin (total) and for tyrosinated tubulin and acetylated tubulin. Solid yellow contour in the overlay marks the area used for detection of individual filaments. Scale bar 5 µm. Middle row: Zoom-ins corresponding to the dashed yellow square in the top row. Bottom row: Example of individual filament detection in each channel (first three panels) and binarized overlay of the detections (right panel). (C) Schematics of individual filament analysis: detection was performed in each channel separately using maximum intensity projection. For each detected segment, the intensity was quantified in all three channels. (D) Outlines of microtubule filaments detected and quantified in the total tubulin channel (for the cell depicted in (A)), color-coded according to their background corrected integrated intensity. (E) Histogram of the background-corrected integrated intensity of individual filaments detected in all three channels and quantified in total tubulin channel for the cell shown in (A) (black dots, n=1736). The solid black line shows the fit of sum of two Gaussian functions: the first corresponds to a single filament intensity distribution (dashed blue line) and the second Gaussian corresponds to the double filament intensity distribution, i.e. first Gaussian convoluted with itself (dashed orange line). Dashed lines mark the mean and mean plus standard deviation of the first Gaussian. The latter was used as a threshold for single microtubule filtering. (F) Illustration of single filament intensity filtering: outlines of the filaments with intensity below the threshold are colored in blue (assigned as a single microtubule) and filaments above it in orange (assigned as two or more microtubule bundles). (G) Two color heatmap overlay of normalized intensity distributions of single microtubule segments detected in tyrosinated (cyan, n=10281 segments) and acetylated (magenta, n=9369 segments) channels and quantified in both (22 cells, N=2 independent experiments). (H) Average normalized level of tyrosination per cell for single microtubule segments detected in the acetylated channel (*α*, magenta) and average normalized level of acetylation for segments detected in tyrosinated channel (*β*, cyan). Horizontal black lines mark mean ± S.E.M. (22 cells, N=2 independent experiments). (I) Illustration of analysis pipeline for a quantification of tyrosinated and acetylated microtubules number in dendrites.

As a first step we performed three-color STED microscopy in the soma and dendrites of DIV9 cells to detect total (alpha-)tubulin, tyrosinated tubulin, and acetylated tubulin. For the analysis of singe-microtubule intensities we used a subvolume of the cell body just below the nucleus (Fig.2A,B), where the majority of microtubules were located in the *x,y* plane and confined to a relatively thin flat layer. We established a custom curvilinear structure detection algorithm to detect filament segments in all three channels and to quantify their background-corrected fluorescence intensity for all channels (Fig.2B-D, S2A) [23].

Next, we focused on the robust estimation of average single filament intensity in the total tubulin channel. We observed that the average intensity of total tubulin was slightly lower for segments detected using acetylated tubulin, compared to segments detected using either total and tyrosinated tubulin (Fig.S2B). This could be caused by a subset of highly modified microtubules that are stained less effectively by the alpha-tubulin antibody. To take this into account, we pooled together the total tubulin intensities of all segments, independent of the detection channel (Fig.2E). This histogram displayed a skewed distribution with a tail in a range of higher intensities (Fig.2E). We reasoned that the peak of the distribution represents the intensities of single microtubules, while the tail corresponds to the presence of microtubule bundles containing two or more overlapping microtubules, as both of these groups could clearly be distinguished in the original images (Fig.2B,D). To obtain a robust estimate for the intensity of individual microtubules, we fitted the histograms with a sum of two Gaussian distributions (Fig.2E). The first Gaussian represents the intensity distribution of a single microtubule, with a standard deviation determined by several different factors (antibody staining variations, microtubules going in and out of focus). The second Gaussian corresponds to bundles of two microtubules and mathematically represents a convolution of first Gaussian with itself (assuming random independent intensity sampling of two microtubules in a bundle). We used the mean value of the first Gaussian as an estimate of average single microtubule intensity in the total channel.

We proceeded with an estimation of the average levels of tyrosination and acetylation per individual microtubule (per cell). To exclude segments corresponding to the bundles of multiple microtubules, we only analyzed segments for which the total tubulin intensity was below the mean intensity of the first Gaussian plus one standard deviation (Fig.2E). Visual inspection of segments below and above the threshold confirmed that this filtering eliminated the majority of thick or bright bundles (Fig.2F). Consistently, including only the single-MT segments determined from total tubulin intensities also resulted in more unimodal and symmetric distributions for the intensities of tyrosination and acetylation levels of segments identified in these respective channels (Fig.S2C, Fig.2G). The average intensities of single tyrosinated or acetylated microtubule were estimated as the average values of intensities detected and quantified in the same corresponding channel. These values were used for the normalization of intensities shown at Fig. 2G and S2D,E. In addition, we also quantified the levels of tyrosination and acetylation of all segments detected in the acetylation and tyrosination channel, respectively. This analysis enabled us to build two two-dimensional histograms that show the levels of both tyrosination and acetylation for microtubule segments detected either in the acetylation channel or the tyrosination channel (Fig 2G).

The resulting histograms show that segments detected by acetylation have, on average, lower levels of tyrosination than segments detected by tyrosination, and vice versa (Fig. 2G). This quantitatively confirms the general impression that these chemical groups mark two different subsets and that microtubules with high levels of acetylation are mostly detyrosinated. However, despite clearly separating into two subsets, even highly acetylated microtubules display residual tyrosination, whereas many tyrosinated microtubules have some extent of acetylation. The measured relative level of tyrosination for acetylated microtubules, compared to average tyrosination of tyrosinated microtubules, which we termed *α*, was equal to 0.53 ± 0.11 (average ± SD) and the level of acetylation for tyrosinated microtubules, compared to the acetylation of acetylated microtubules, termed *β*, was 0.45 ± 0.07 (Fig.2H). These data demonstrate that microtubules can be divided in two different subsets based on the detection of tyrosination and acetylation. We will refer to microtubules detected in the acetylated tubulin channel, displaying on average 47% lower levels of tyrosination than microtubules detected in the tyrosinated channel, as stable microtubules. Likewise, dynamic microtubules are identified as microtubules detected in the tyrosinated tubulin channel and feature 55% lower levels of acetylation than microtubules detected in the acetylated channel.

We next set out to use the intensities of total tubulin, acetylation and tyrosination on individual microtubules to determine the both the total number of microtubules within dendrites, as well as the number of stable and dynamic subsets within dendrites. To estimate the total number of microtubules, the dendritic intensity of total tubulin could just be divided by the single-microtubule intensity (assuming consistent labeling throughout the neuron). However, for the quantification of stable and dynamic microtubules, we needed to account for the “chemical crosstalk” that we observed, i.e. the tyrosination and acetylation levels detected for stable and dynamic microtubules, respectively. As a result, the integrated tyrosinated intensity of a dendrite was not just the sum of intensities of dynamic microtubules, but also included the contribution from the residual tyrosination of stable microtubules (and vice versa). This situation was analogous to instances of spectral crosstalk in fluorescent microscopy, where emission from one dye is detected in the spectral channel of another dye [24], and we therefore used standard formulas for spectral unmixing and our estimates for *a* and *β* (Fig.2H,I) to take this posttranslational modification crosstalk into account.

To calculate the composition of the dendritic microtubule network, we focused on the proximal 5-10 μm of a dendrite (Fig.3A,B) and used the corresponding single-filament intensity and crosstalk estimates from the soma of the same cell. First of all, this showed that the total number of microtubules in a dendrite depended linearly on its cross-sectional area in the range from 1 to 10 μm^2^ with a slope of 68 microtubules per μm^2^. In addition, it revealed that dendrites have more than four times more acetylated microtubules than tyrosinated microtubules (74 ± 8% versus 16 ± 11%, average ± SD) and that this factor was largely independent of the diameter of the dendrite (Fig.3C,D,E). We furthermore found that these two subsets did not completely account for the total number of microtubules that we measured, leaving a small fraction of 10 ± 14% of microtubules that were neither acetylated, nor tyrosinated.

A potential weakness of the analysis that we performed is that it assumes that the levels of acetylation and tyrosination on stable and dynamic subsets measured in the cell body are comparable to those within dendrites. To overcome this, single-microtubule levels of acetylation and tyrosination would need to be measured directly from the dendrites, which requires 3D images in which single dendritic microtubules are clearly distinguished. Because this was not possible using our STED microscopy approach, we switched to expansion microscopy (ExM) to improve both lateral and axial resolution [10, 25]. In expansion microscopy, stained samples are embedded in and crosslinked to a swellable hydrogel, followed by proteolytic digestion and physical expansion, which will increase the spacing between the remaining gel-linked protein fractions and fluorophores. Since gels expand in all dimensions, this leads to an isotropic resolution improvement determined by the expansion factor of the gel (about 4 times).

Indeed, expanded samples demonstrated a substantial increase in the clarity with which microtubule organization could be perceived (Fig.4A,B,S3A, Supplementary Movie 2). We therefore repeated our analysis of the spatial distribution of tyrosinated and acetylated microtubules and found that the peripheral enrichment of tyrosinated microtubules was even more pronounced in ExM samples, as shown in *y,z* cross-section images (Fig.4B) and radial distribution plots (Fig.4C,D,E,F). Even though visual tracing of individual filaments remained challenging (Fig.S3A), we were able to estimate the relative abundance of acetylated and tyrosinated microtubules by decomposing the radial density of total tubulin as a sum of the acetylated and tyrosinated radial densities (Fig.4G). Although this analysis does not take into account the fraction of microtubules that is neither tyrosinated or acetylated, it independently confirms the prevalence of acetylated microtubules (65%) over tyrosinated (35%).

The successful decomposition of total tubulin using only these two subsets, as well as the higher fraction of tyrosinated tubulin in comparison with our estimate from soma-based intensity rescaling (Fig. 3E), prompted us repeat our microtubule counting using dendrite-based intensity rescaling. Unfortunately, our ExM data also did not have sufficient resolution to resolve enough individual microtubules to reliably determine the single-microtubule estimates of acetylation and tyrosination required for such analysis. First, we tried to use ExSTED microscopy to improve resolution [26] and found that three-color volumetric STED acquisition of ExM samples resulted in substantial photobleaching, which strongly impaired the integrated intensity analysis (data not shown). We then realized that most dendritic microtubules are organized in bundles that run parallel to the coverslip and that resolving microtubules would be easier if we could alter the sample orientation such that microtubules are aligned with the optical axis of our microscope. Since in a regular ExM acquisition the axial (the poorest) dimension of PSF is oriented perpendicular to the filaments (located parallel to the coverslip plane), we decided to generate thick gel slices that were rotated by 90 degrees, a procedure we termed FlipExM (Fig.5A,B). In this configuration we exploit the better lateral resolution to resolve microtubules, while PSF blurring along the optical axis happens parallel to filaments (Fig.5B, Supplementary Movies 3 and 4).

**Figure 3.**
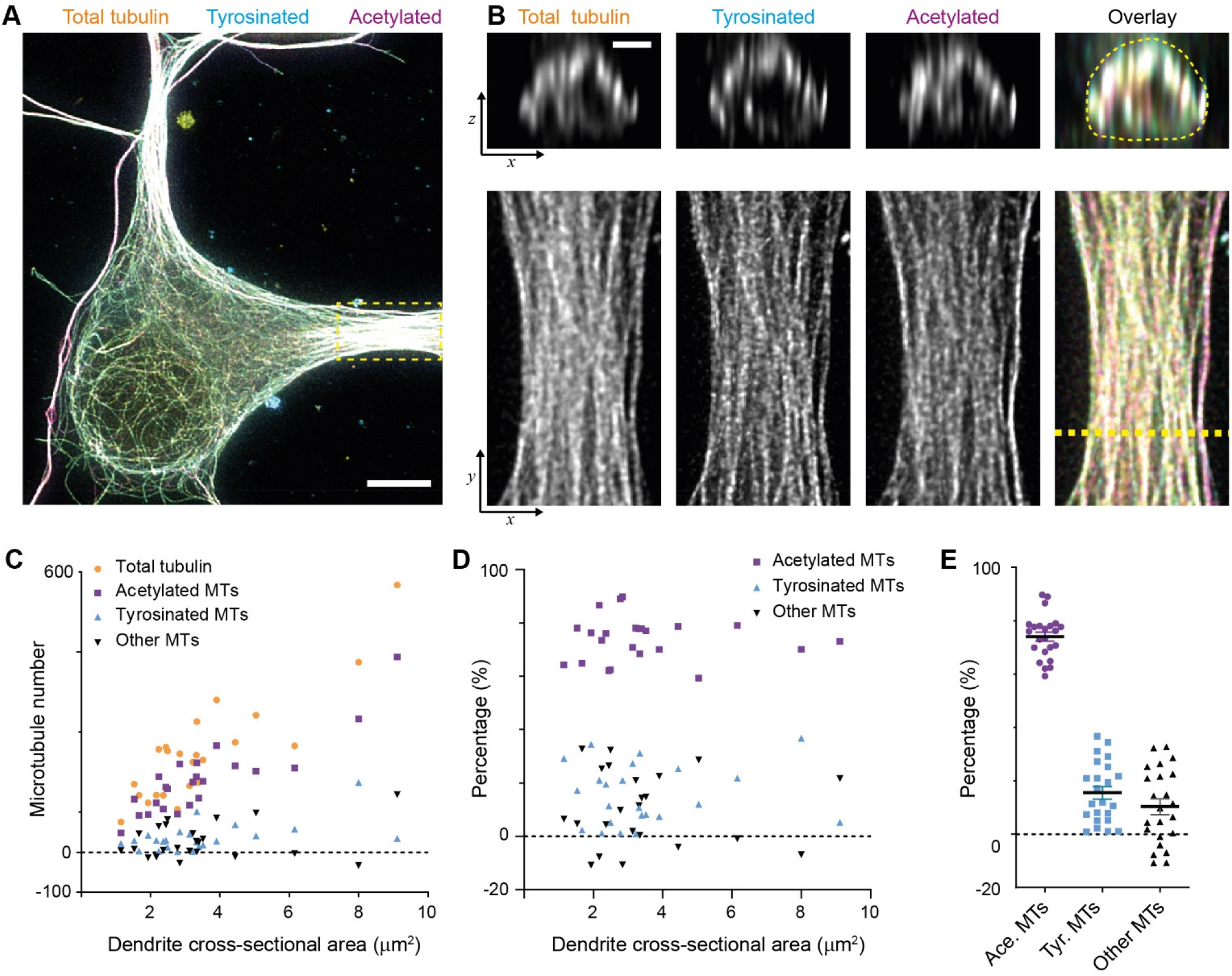
Estimation of absolute numbers of (modified) microtubules in dendrites using single-microtubule intensities from the soma. (A) Three-color overlay of maximum intensity projection of a 2D STED z-stack including the whole volume of dendrites (total tubulin in yellow, tyrosinated in cyan and acetylated in magenta). Dashed yellow rectangle marks zoom-in shown in (B). Scale bar 10 µm. (B).Maximum intensity projections of a segment of dendrite (bottom row, marked by yellow dashed rectangle in (A)) and individual YZ slices (top row, corresponds to a dashed yellow line). (C)Numbers of total, tyrosinated, acetylated and other (i.e. neither tyrosinated nor acetylated) microtubules per dendrite as a function of cross-sectional area (n=23 dendrites, N=2 independent experiments). These numbers were determined using the approach outlined in Figure 1. (D)-(E) Percentage of tyrosinated, acetylated and other microtubules per dendrite as a function of dendrite’s cross section area (D) or pooled together (E). Horizontal black lines in (E) mark mean ± S.E.M. (n=23 dendrites, N=2 independent experiments).

**Figure 5.**
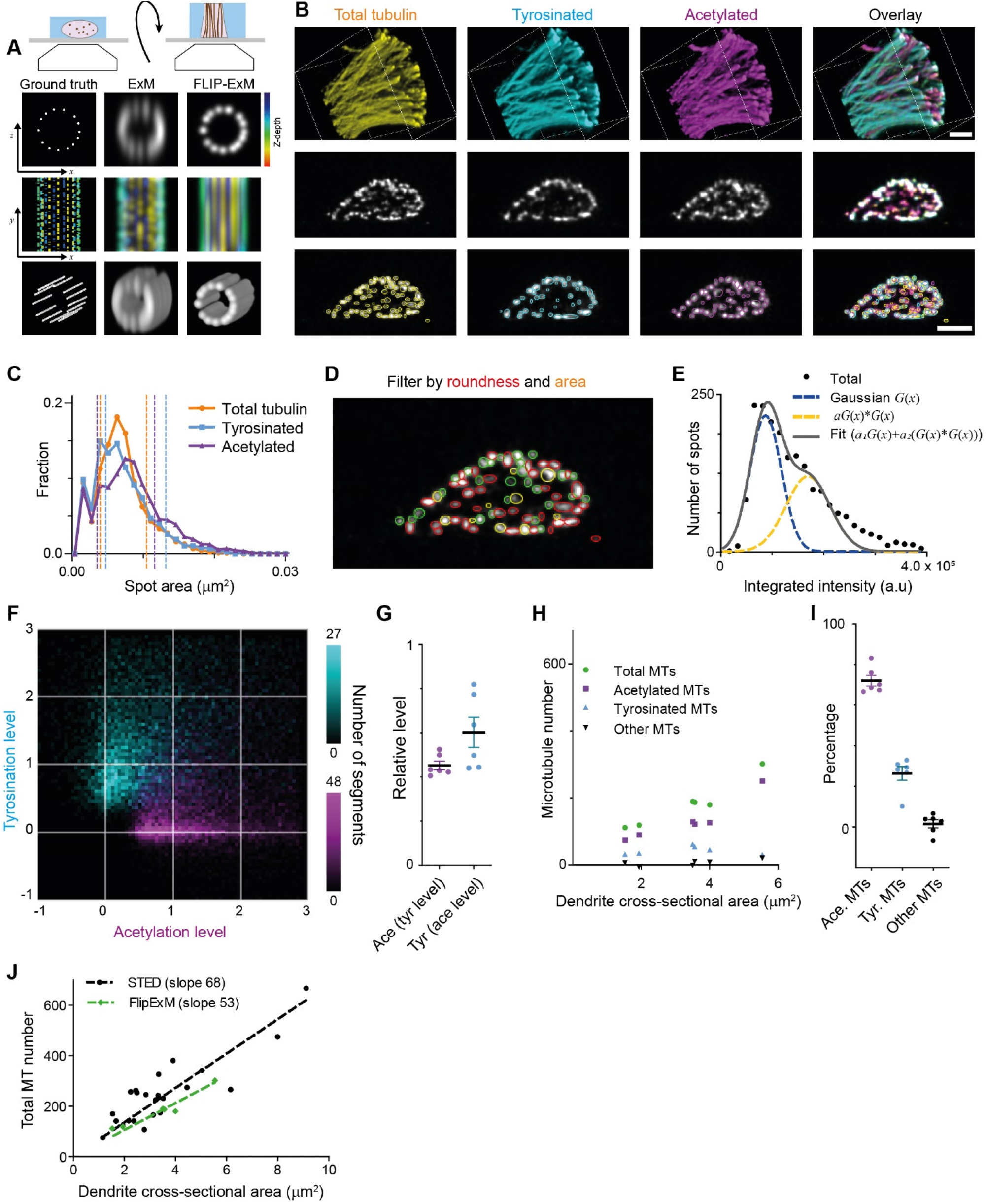
Direct estimation of microtubule numbers in dendrites using FlipExM. (A) Top: Schematics of gel reorientation for FlipExM imaging. Bottom: Simulated z-stacks illustrating the advantages of FlipExM for imaging of dendritic microtubules. A set of simulated circumferential microtubules in a dendrite of 1 µm diameter (left) were convoluted with a point spread function corresponding to a regular ExM (middle) or FlipExM (right) imaging (top to bottom: single XZ plane, color-coded depth projection, 3D rendering). (B) Representative volumetric 3D rendering (top) and single YZ slices (middle) of total tubulin and its posttranslational modifications in a dendrite imaged using FlipExM. The bottom row shows automatic detections of microtubules in cross sections. Scale bars 1 μm (physical size post-expansion 4.15 μm). (C) Area histogram of spots corresponding to microtubules cross sections in three channels for the dendrite shown in (B). Spots were pre-filtered using roundness criteria (n=7103, 4844, 3821 for total, tyrosinated and acetylated channels). An interval between dashed lines marks the range for spot’s area filtering. Abscissa units are recalculated according to expansion factor (4.15). (D) Single YZ slice of total tubulin channel with detections filtered by roundness marked by red circles, detections filtered by area marked by yellow circles and remaining detections attributed to single microtubules marked in green. (E) Histogram of background subtracted integrated intensity of individual microtubules cross sections detected and quantified in total tubulin channel for the dendrite shown in (B) (black dots, n=1981). The solid black line shows the fit of sum of two Gaussian functions: the first corresponds to a single microtubule cross section intensity distribution (dashed green line) and the second Gaussian corresponds to the double cross section intensity distribution, i.e. first Gaussian convoluted with itself (dashed red line). (F) Two color heatmap overlay of normalized intensity distributions of single microtubule cross sections detected in tyrosinated (cyan, n=8642 spots) and acetylated (magenta, n=12552 spots) channels and quantified in both (6 cells, N=2 independent experiments). (G) Average normalized level of tyrosination per cell for single microtubule cross section detected in acetylated channel (*α*, magenta) and average normalized level of acetylation for cross sections detected in tyrosinated channel (*β*, cyan). Horizontal black lines mark mean ± S.E.M. (6 cells, N=2 independent experiments). (H) Numbers of total, tyrosinated, acetylated and non-modified microtubules per dendrite depending on dendrite’s cross section area (n=6 cells, N=2 independent experiments). (I) Percentage of tyrosinated, acetylated and non-modified microtubules as a fraction of total microtubule number per dendrite. Horizontal black lines mark mean ± S.E.M. (n=6 cells, N=2 independent experiments). (J) Number of microtubules per dendrite determined using STED (black dots, data from Fig.2C) or FlipExM (green dots, data from (H)) imaging. Dashed lines show independent linear fits passing through the origin.

Although we still could not discern individual microtubules within tight bundles, we observed many individual microtubules traversing through the dendritic volume. We therefore set out to quantify the intensities of these microtubules, so that these could be used to quantify the total number of microtubules and the abundance of microtubule subsets. The cross-section of individual filaments in dendrite’s cross-sections were automatically detected in each channel (Fig.5B, bottom row), and we quantified their area and background-corrected intensity in each channel. To exclude noise and bundles, we then applied area and roundness filters on our detections (Fig.5C,D). After this geometrical filtering, the intensity distribution showed a similar bimodal or skewed shape as found earlier for the filaments in the cell body (Fig.5E, Fig.2E, S4A,B). Therefore, we again used curve fitting (similar to Fig.2) to estimate the average intensity of microtubule cross-sections in each channel. The distributions of tyrosinated and acetylated detections in the tyrosination/acetylation plane (Fig.5F, S4C,D) appeared very similar to the data obtained earlier using the cell body (Fig.2G), but with more distinct separation between the two clusters. Compared to the cell body data, we found slightly different values for the average tyrosination level of acetylated microtubules (0.45 ± 0.05), as well as the acetylation level for tyrosinated (0.60 ± 0.17) (Fig.5G).

Finally, we used the single-microtubule intensity levels measured directly within dendrites to quantify total microtubule numbers, as well as the number of acetylated and tyrosinated microtubules (Fig. 5G-J). First, we divided the integrated cross-section intensity of the total tubulin channel by our single cross-section intensity estimate for dendrites with different diameters (Fig. 5G-J). A linear fit through the total number of microtubules as a function of cross-sectional area yielding an estimated microtubule density of 68 and 53 microtubules per square micrometer for the cell body and dendrite methods, respectively (compared to 68 microtubules per square micrometer for the cell body method, Fig.5J). Next, to determine the number of acetylated and tyrosinated microtubules, we employed the “modification unmixing” approach mentioned previously. Consistent with our earlier results, this analysis revealed that stable, acetylated microtubules formed the largest population (72 ± 6%). The fraction of tyrosinated microtubules was larger than our earlier estimate (26 ± 8%), at the expense of the fraction of microtubules that were neither acetylated nor tyrosinated (2 ± 5%, Fig.5H,I, S4E). These results indicate that acetylated and tyrosinated microtubules together account for 98% of all dendritic microtubules, with acetylated microtubules being almost three times more abundant.

## DISCUSSION

The high density of the neuronal microtubule cytoskeleton has so far obscured its exact composition and organization. Earlier work has used electron microscopy to reveal the number and spatial organization in dendrite cross-sections, but this technology is difficult to combine with the robust detection of distinct subsets [21, 22]. Here we used super-resolution light microscopy to explore the composition and architecture of the microtubule cytoskeleton in dendrites. In addition to visualizing all microtubules using an antibody against alpha-tubulin, we focused on two microtubule subsets: those labeled using antibodies against acetylation and tyrosination, typically classified as stable and dynamic microtubules [13, 16, 27]. Volumetric STED and expansion microscopy revealed a striking spatial organization in which stable, acetylated microtubules are enriched in the core of the dendritic shaft, surrounded by a shell of dynamic, tyrosinated microtubules (Fig. 1, 4). While our earlier two-dimensional super-resolution imaging already hinted at a spatial separation between different subsets [17], the current work provides the first quantitative three-dimensional mapping of subset organization throughout a large set of dendrites.

**Figure 4.**
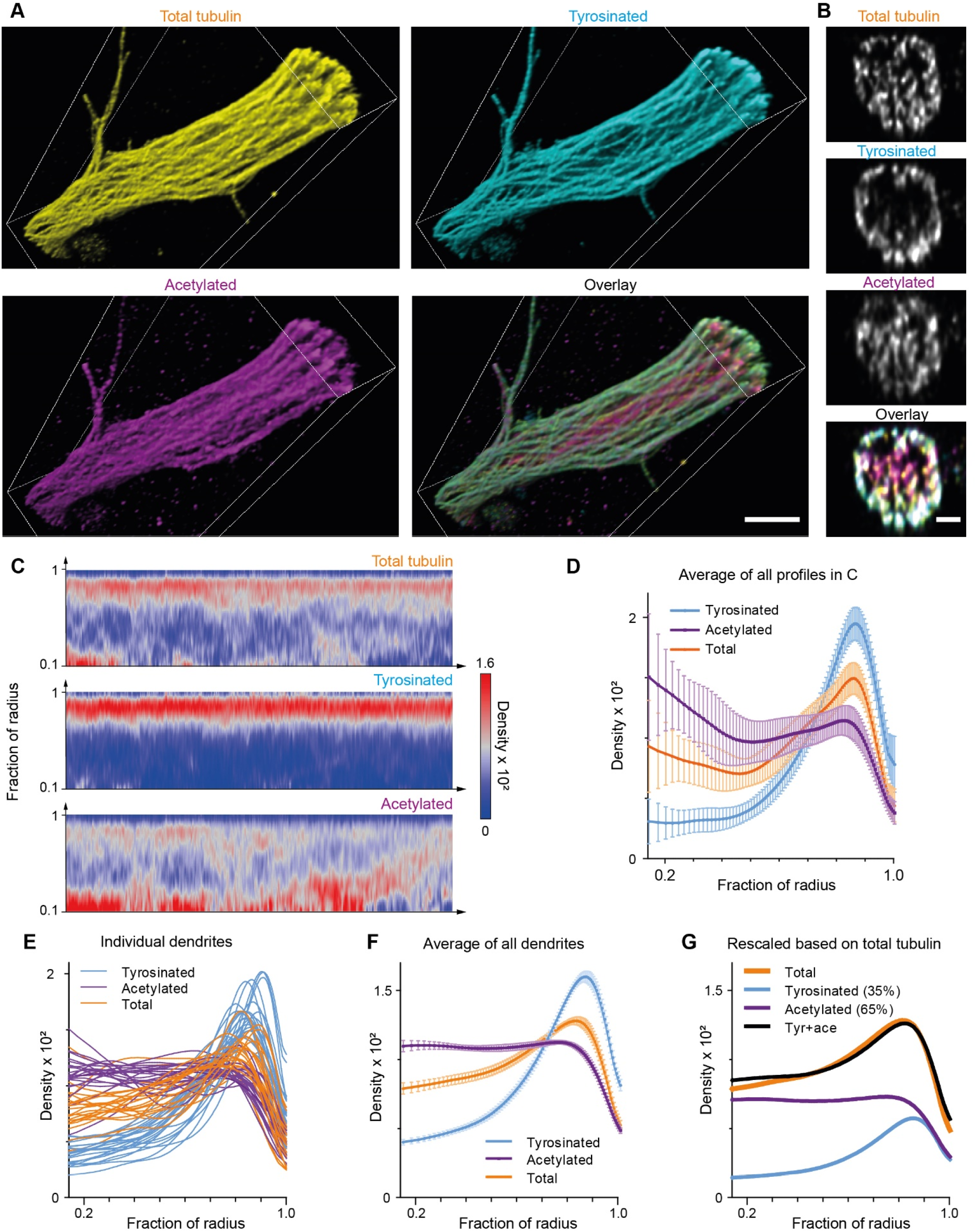
Expansion Microscopy improves quantification of the radial distribution of microtubule modifications in dendrites. (A) Representative volumetric 3D rendering of total tubulin and its posttranslational modifications in a dendrite imaged using ExM. Scale bar 2 μm (physical size post-expansion 8.3 μm). (B) Representative single YZ cross section of the dendrite from (A). Scale bar 0.5 μm (physical size post-expansion 2.07 μm). (C) Heatmaps of normalized radial intensity distributions for total tubulin (top), tyrosinated (middle) and acetylated (bottom) microtubule posttranslational modifications along the length of the dendrite shown in (A). Abscissa units are recalculated according to expansion factor (17 µm equals to 70.5 µm physical size post-expansion). (D) Radial distribution of total tubulin (orange), tyrosinated (cyan) and acetylated (magenta) microtubule posttranslational modifications averaged over the length of the dendrite shown in (A) (n=404 profiles). Error bars represent SD. (E)-(F) Radial distribution of total tubulin (orange) and tyrosinated (cyan) and acetylated (magenta) posttranslational modifications intensities averaged per dendrite (E) and among all dendrites (F) imaged using ExM (n=9460 profiles, 22 cells, N=2 independent experiments). Error bars represent S.E.M. (G) Decomposition of total tubulin radial intensity distribution as a weighted sum of tyrosinated and acetylated distributions.

The enrichment of dynamic microtubules near the plasma membrane is consistent with the well-established interplay between growing microtubule plus ends and (sub)cortical complexes [28, 29]. More specifically, dynamic microtubules have been shown to regularly invade into dendritic spines to facilitate intracellular transport or regulate spine morphology in response of specific synaptic stimuli [30-34]. Next to ensuring the enrichment of dynamic microtubules near the plasma membrane, the spatial separation between stable and dynamic microtubules might also promote efficient intracellular transport by separating cargoes driven by subset-specific motors. Moreover, for motors that do not discriminate between microtubule subsets, the spatial separation between mostly minus-end out oriented stable microtubules and mostly plus-end out oriented dynamic microtubules will facilitate directional transport by limiting directional switching induced by cargo-attached motors binding to neighboring microtubules of opposite polarity. In future work, we will explore how the transport patterns of different cargoes depend on the associated motors and the organization of the neuronal microtubule cytoskeleton.

The use of three-color super-resolution imaging allowed us to include a marker for total tubulin and, in combination with novel analysis methods, provide two independent estimates for the total number of microtubules in dendrite sections, as well as the number of acetylated and tyrosinated microtubules (Fig. 3, 5). Our estimates for the total microtubules were obtained by dividing the total intensity of a generic tubulin stain by the intensity measured on individual microtubules in either the soma (STED, Fig. 3) or dendrite itself (Flip-ExM, Fig. 5), which revealed an average density of 68 or 53 microtubules per μm^2^, respectively. These values are consistent with earlier estimates using electron microscopy (66 microtubules per μm^2^)[22]. Although we used various filtering steps to prevent mistaking small microtubule bundles for individual microtubules, it remains possible that occasional inclusion of such bundles increased our estimate for single microtubules, thereby lowering our estimate for the total number of microtubules. Alternatively, these differences could be caused by local differences in expansion factor or just reflect sample-to-sample differences in the number of microtubules per area. Nonetheless, the close correspondence between our estimates and values obtained using electron microscopy on dendritic cross-sections demonstrates the strength of combining super-resolution microscopy with quantitative analysis.

In all our measurements, acetylated microtubules strongly outnumber tyrosinated microtubules. The two single-microtubule calibration methods (i.e. soma versus dendrite) yielded strikingly similar estimates for the percentage of acetylated microtubules (74 ± 8% versus 72 ± 6%), while their estimates for tyrosinated and other (non-tyrosinated and non-acetylated) microtubules differed to some extent (16% tyrosinated and 10% other microtubules versus 26% and 2% for soma and dendrite estimations, respectively). These results suggest that the acetylation level of stable microtubules is similar between soma and dendrites, whereas the tyrosination level of dynamic microtubules could be higher in the soma than the dendrite. For the soma-based method, this would overestimate the dendritic single-microtubule tyrosination levels and result in undercounting dendritic tyrosinated microtubules, leaving a larger fraction of *other* microtubules. The idea that dendritic dynamic microtubules are on average more detyrosinated is consistent with our finding that these microtubules also have higher levels of acetylation compared to the soma (Fig. 5G). We therefore consider the dendrite-intrinsic measurements to be more reliable, i.e. 72 ± 6% acetylated, 26 ± 8% tyrosinated and 2 ± 5% other microtubules.

It is important to note that the two markers that we used, acetylation and tyrosination, represent only a small part of the possibly ways in the microtubule surface can become differentiated, such as through other modifications like polyglutamylation, phosphorylation, palmitoylation, incorporation of different tubulin isoforms, and adsorption of different MAPs [9, 13]. Remarkably, our analyses nonetheless revealed that labeling acetylated and tyrosinated microtubules leaves only a very small fraction (2%) of microtubules unlabeled. This suggests that most detyrosinated microtubules in dendrites are acetylated and that most other modifications or MAPs are found on microtubules that are tyrosinated or acetylated.

In this work, we have introduced innovative imaging and analysis approaches to quantitatively map the neuronal cytoskeleton. In future work, we aim to map how other modifications and various microtubule-associated proteins are distributed over these two subsets of microtubules. In addition, the distribution of modifications and microtubule-associated proteins along the length of individual microtubules should be mapped to better understand how dynamic microtubules may become stabilized. We anticipate that such experiments will benefit from ongoing advances in expansion microscopy, such as iterative expansion or single step approaches with higher expansion factors.

## Supporting information

Vidoe 4

## ACKNOWLEDGEMENTS

This work was supported by the European Research Council (ERC Consolidator Grant 819219) and ZonMW (project 91217002).

## DATA AVAILABILITY

The data that support the findings of this study (graphs including raw data, ImageJ macros and Matlab code used for analysis) are freely available online at the ‘figshare’ repository https://doi.org/10.6084/m9.figshare.c.5306546.v1. Links to the source code of used ImageJ plugins and their archived versions are provided in the text. Raw microscopy images are available on request.

## MATERIAL AND METHODS

### Primary rat neuron culture and immunostaining

Dissociated hippocampal neuron cultures were prepared from embryonic day 18 rat pups of mixed gender according to the previously published protocol [35]. Briefly, cells were plated on 18-mm glass coverslips coated with poly-l-lysine (37.5 mg/ml) and laminin (1.25 mg/ml) in a 12-well plate at a density of 50k/well. Cultures were maintained in Neurobasal medium (NB) supplemented with 2% B27, 0.5 mM glutamine, 16.6 μM glutamate, and 1% penicillin/streptomycin at 37°C in 5% CO2.

We followed our recently published immunostaining/expansion protocol, described in details in [10]. In short, at DIV9 neurons were pre-extracted for 1 minute using 500 µl of 0.3% Triton X-100 (Sigma X100), 0.1% glutaraldehyde (Sigma G7526) in MRB80 buffer (80 mM Pipes (Sigma P1851), 1 mM EGTA (Sigma E4378), 4 mM MgCl_2_, pH 6.8) and fixed for 10 minutes using 4% PFA (Electron Microscopy Sciences 15,710) and 4% sucrose in BRB80 buffer (both solutions were pre-warmed to 37 °C). Fixed neurons were washed 3 times in PBS and permeabilized for 10 minute in 500 µl of 0.25% Triton X-100 in BRB80 buffer. Samples were further incubated in 500 µl of blocking buffer (3% w/v BSA in BRB80 buffer) for at least 45 minutes at room temperature. Finally, fixed neurons were sequentially incubated for two hours at room temperature (or overnight at 4 °C) with primary and secondary antibodies diluted in blocking buffer (3% w/v BSA in BRB80 buffer) and washed 3 times in PBS. We used following combinations of primary (dilution 1:200) and secondary (dilution 1:250) antibodies: mouse monoclonal anti-acetylated tubulin (Sigma, [6-11B-1], T7451) with Abberior Star 635P goat anti-mouse IgG (H + L) (Abberior GmbH ST635P-1001-500UG), rat monoclonal anti-tyrosinated tubulin (Abcam, [YL1/2], ab6160) with Alexa Fluor 594 goat anti-rat IgG (H + L) (Molecular Probes, Life Technologies A11007) and rabbit recombinant anti-alpha tubulin antibody (Abcam, [EP1332Y], 52866) with Alexa Fluor 488 goat anti-rabbit IgG (H+L) (Thermo Fisher Scientific, A-11034).

### Expansion microscopy

Expansion microscopy (ExM) was performed according to the proExM protocol [25] with the detailed description published in [10]. Briefly, immunostained neurons on 18-mm glass coverslips were incubated overnight in PBS with 0.1 mg/ml Acryloyl-X (Thermo Fisher, A20770) and afterwards washed three times with PBS. Per coverslip, we made 200 μl of gelation solution by mixing 188 μl of monomer stock solution (1 × PBS, 2 M NaCl, 8.625% (w/w) sodium acrylate (SA)(Sigma Aldrich 408220), 2.5% (w/w) acrylamide (AA), 0.15% (w/w) N,N′-methylenebisacrylamide), 8 μl of 10% (w/w) tetramethylethylenediamine (TEMED, BioRad 161-0800) accelerator and 4 μl of 10% (w/w) ammonium persulfate (APS, Sigma Aldrich A3678) initiator (added at the last step). 120 μl of gelation solution was transferred to a gelation chamber, made of silicone mold with inner diameter of 13 mm (Sigma-Aldrich, GBL664107) attached to a parafilm-covered glass slide. The sample was put cell-down on top of the chamber to close it off. After incubation at RT for 1-3 min, the sample was transferred to a humidified 37°C incubator for at least 30 min to fully polymerize the gel. After gelation, the gel was transferred to a 12-well plate with 2 ml of digestion buffer (1 × TAE buffer (40 mM Tris, 20 mM acetic acid, 1 mM EDTA, pH8), 0.5% Triton X-100, 800 mM NaCl, 8 U/mL proteinase-K (ThermoFisher, EO0492)) for 4 h at 37°C for digestion. The gel was transferred to 50 ml deionized water for overnight expansion, and water was refreshed once to ensure the expansion reached plateau. Plasma-cleaned 24 × 50 mm rectangular coverslips (VWR 631-0146) for gel imaging were incubated with 0.1% poly-l-lysine to reduce drift of the gel during acquisition. The gel was mounted using custom-printed imaging chambers [10]. The expansion factor was calculated for each sample as a ratio of a gel’s diameter to the diameter of the gelation chamber and was in the range of 4.14–4.16. For the FlipExM samples, we cut a thin piece of gel (1 cm x 3 mm) using a razor blade and flipped it on its cut edge during transfer to the imaging chamber.

### STED imaging

Data from non-expanded samples were acquired using a Leica TCS SP8 STED 3X microscope with a pulsed (80MHz) white-light laser, HyD detectors and spectroscopic detection using HC PL APO 100×/1.40 Oil STED WHITE (Leica 15506378) oil-immersion objective. For Abberior STAR 635P and Alexa 594 we used 633 nm and 594 nm laser lines for excitation and a 775 nm synchronized pulsed laser for depletion, with a time gating range of 0.3-7 ns. For Alexa 488 we used 488 nm excitation, 592 nm continuous depletion laser line and time gate of 1.1-7 ns. Emission detection windows were 500-560 nm, 605-630 nm and 640-750 nm for Alexa 488, Alexa 594 and Abberior STAR 635P, respectively. No bleed-through was observed between the channels. For three-color cell body imaging (Fig.2-3), each fluorescent channel was imaged using the 2D STED configuration (vortex phase mask) in sequential z-stack mode from highest to lower wavelength, to prevent photobleaching by the 592 nm depletion laser line. For two-color imaging of dendrites (Fig.1) we used the Abberior STAR 635P/Alexa 594 combination and a single 775 nm depletion line and therefore acquired images in line-sequential mode. For the 3D STED imaging, we used a combined depletion PSF light path consisting of a mixture 60% Z-donut and 40% vortex phase mask, providing approximately isotropic resolution.

For the data shown in Fig.2-3, the size of the field-of-view was in the range of 30-50 µm and it was positioned to include the whole cell body of a neuron (soma) and the first 5-10 µm of dendrites emanating from it (Fig.2A,3A). For the data shown in Fig.1, the size of the field-of-view was in the range of 50-100 µm and it covered 30-50 µm of the proximal dendrites. The depth of z-stacks varied in the range from 3 to 6 µm for each acquisition and for all cases it was chosen to fully cover the dendrite’s thickness. Lateral pixel size was in the range of 27-30 nm with a distance between z-planes in the range of 150-160 nm. The z-stacks were subjected to a mild deconvolution using Huygens Professional software version 17.04 (Scientific Volume Imaging, The Netherlands) with CMLE (classic maximum likelihood estimation) algorithm with parameters of SNR (Signal-to-Noise Ratio) equal to 7 over 10 iterations. After the deconvolution, z-stacks of tyrosinated and acetylated channels were registered in 3D to total tubulin channel using maximum intensity projections in XY and XZ planes using Correlescence plugin v.0.0.4 (https://github.com/ekatrukha/Correlescence archived on Zenodo repository https://doi.org/10.5281/zenodo.4534715) for ImageJ.

### ExM/FlipExM samples imaging

Expanded gels were imaged using the same Leica TCS SP8 STED 3X microscope with a pulsed (80 MHz) white-light laser, HyD detectors and spectroscopic detection using HC PL APO 86 ×/1.20 W motCORR STED (Leica 15506333) water-immersion objective with corrective collar. Each fluorescent channel was imaged in confocal line-sequential mode. For Alexa488 we used 488 nm excitation and 500-560 nm emission range, for Alexa594 we used 594 nm excitation and 605-630 nm emission and Abberior STAR 635P we used 633nm excitation and 640-750 nm emission. For ExM samples, the size of field-of-view was in the range of 50-100 µm and thickness in the range of 10-20 µm, chosen to cover the whole volume of a dendrite. Dimensions of FlipExM stacks were 20-30 µm in XY and 30-50 µm in Z. In both cases the pixel size in XY plane was in the range of 60-80 nm and the distance between z-planes was in the range of 150-180 nm. The z-stacks were subjected to a mild deconvolution using Huygens Professional software version 17.10 (Scientific Volume Imaging, The Netherlands) with CMLE (classic maximum likelihood estimation) algorithm with parameters of SNR (Signal-to-Noise Ratio) equal to 15 over 10 iterations.

### Single microtubule intensity estimate in the cell body using STED

From registered z-stacks we chose substacks of 4-6 frames (0.7-1 µm thick) located at the bottom of the cell under the nucleus (Fig.2A), where the density of microtubule network was low. Using maximum intensity projections of these substacks in each fluorescent channel, we extracted segments of microtubule filaments using CurveTrace plugin ver.0.3.5 (https://github.com/ekatrukha/CurveTrace archived on Zenodo repository https://doi.org/10.5281/zenodo.4534721) for ImageJ implementing [23].

Detection parameters used were: line width of 2.5 pixels (75 nm) (standard deviation of line thickness) and minimum segment’s length of 0.6 µm. The detection of filaments was limited to the area of cell body, excluding dendrites (Fig. 2B, S2A). After detection, each segment of microtubule was stored as a polyline ROI (region of interest) file in ImageJ format, essentially represented as a set of ordered XY coordinates. The detection was performed separately for each fluorescent channel, to take advantage of sparser filament’s subnetworks with less overlap, displayed in tyrosinated and acetylated channel (Fig.2B, bottom row).

The quantification of filament intensities was performed on the sum of slices of the substacks used for the detection (SUM projection). The intensity of filament segments detected in the different channels was quantified for each fluorescent channel (total, tyrosinated, acetylated), producing nine datasets (Fig.2C). For each detected polyline ROI segment of length *L*_mid_ we first measured the integrated intensity *I*_mid_ with a line width *W*_microtubule_ of 6 pixels (∼180 nm) covering the area of *S*_mid_. In this case *W*_microtubule_ corresponds to the width used to normalize intensity of a single microtubule segment. As a second step, we measured the integrated intensity *I*_wide_ of the same segment with a wider line’s width of 13 pixels (∼390 nm) covering the area of *S*_wide_. From these measurements, the average intensity of the background *I*_BG_ was calculated as:

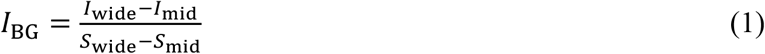

and the background-corrected average intensity of a segment *I*_segm_ as:

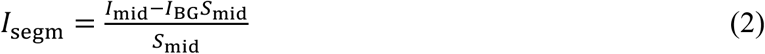

Since the line width was fixed, this value does not depend on the length of the filament and essentially represents average fluorescent intensity along the filament. The described measurements were automated using an ImageJ script [36].

The histogram of segment intensities in total channel (pooled from all three detections) was fitted with a sum of two Gaussians (Fig.2E) expressed as:

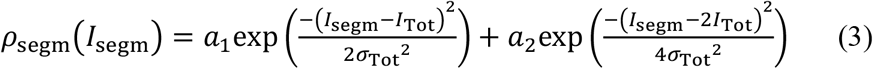

where *a*_1_, *a*_2_ correspond to the amplitudes (weights) of first and second Gaussians, *I*_Tot_ and *σ*_Tot_ are the average intensity and standard deviation of the Gaussian corresponding to the single microtubule intensity distribution (for the second Gaussian, after the convolution, average intensity and standard deviation are 2*I*_Tot_ and 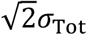) The fitting was performed for each cell individually, to eliminate a difference in imaging conditions and heterogeneity of the sample.

The fitted value of average intensity *I*_Tot_ was used later for the estimation of total microtubule numbers in dendrites (see next section). For quantification of the average levels of tyrosination and acetylation per single microtubule, we introduced a threshold of *I*_Tot_ + *σ*_Tot_ on the corresponding total tubulin intensity of segments detected in acetylated and tyrosinated channels (Fig.2E). Only the segments which total tubulin intensity was below this threshold were used for the calculation of average intensities of single tyrosinated *I*_Tyr_ or acetylated *I*_Ac_ microtubules. The average values for each channel were used for the normalization of intensities presented at Fig.2G,H and S2D,E. The fitting and threshold filtering was performed using custom written MATLAB scripts [36].

### Estimation of microtubules number in dendrites using STED

To estimate the average number of microtubules in dendrites, we first built summary (integrated) XY projection images of z-stacks containing the whole depth of dendrites. Similar to the quantification of single microtubule intensity, we drew a straight line ROI of 2-3 µm (*L*_dendrite_) along a dendrite segment with a width *W*_dendrite_ (using ImageJ). This width varied depending on the dendrite and was chosen to visually include its whole thickness, covering an area of 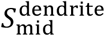. We measured the integrated intensity over this area, denoted as 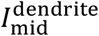 In the second step, we measured the integrated intensity 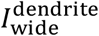 of the same straight line with a width increased by 10 pixels (300 nm) covering the area of 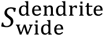 From these measurements, the average intensity of the background 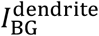 was calculated similar to (1) as:

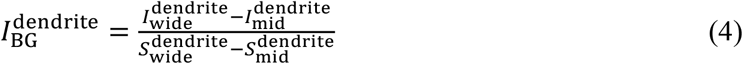

and the background-corrected average intensity per area of a dendrite segment *I*^dendrite^ was calculated as:

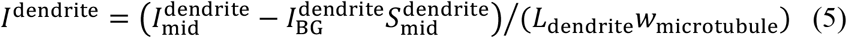

where parameter *W*_microtubule_ is the same width used for estimation of single microtubule segments in the previous section. Expressed in this way, the number of microtubules in total tubulin channel is given by:

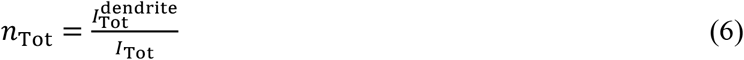

where 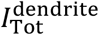 is dendrite’s intensity in total tubulin channel calculated according to (5) and *I*_Tot_ is average single microtubule intensity in total tubulin channel (see previous section). The specific values of *I*_Tot_ were taken from the same cell/z-stack containing the dendrite.

To calculate numbers of microtubules in tyrosinated and acetylated channel we used following formulas (Fig.2I):

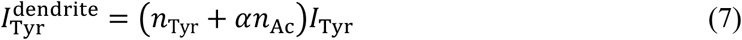

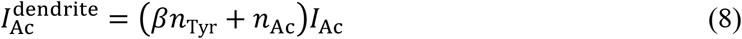

Where 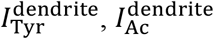 are background corrected dendrite intensities calculated according to (5), *I*_Tyr_ and *I*_Ac_ average single microtubule intensities in tyrosinated and acetylated channel, *α* stands for average level of tyrosination for microtubules detected in the acetylated channel, *β* corresponds to the average acetylation level of microtubules detected in the tyrosinated channel (Fig.2H) and *n*_Tyr_, *n*_Ac_ are numbers of tyrosinated and acetylated microtubules. The solution of system (7)-(8) gives the final formulas:

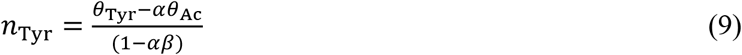

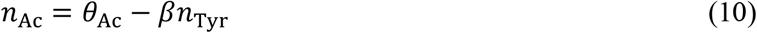

Where 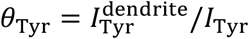 and 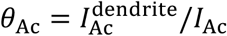. (9)-(10) were used to report the number of tyrosinated and acetylated microtubules in Fig.3C-D. In addition, these values were used to calculate the tyrosinated and acetylated percent of total microtubules number *n*_Tot_ reported in Fig.3E. The number of “non-modified”, other microtubules was calculated as *n*_Other_ = *n*_Tot_ - *n*_Tyr_ - *n*_Ac_.

The dendritic cross section area (Fig.3C-D) was measured by building XZ resliced cross section along a perpendicular line in the area of intensity measurement.

### Radial distribution of intensities in dendrites

We acquired z-stacks covering the whole thickness of a dendrite (using 2D or 3D STED in Fig.1 and confocal for ExM samples in Fig.3). Using maximum intensity projection in XY plane we marked the middle of dendrite with a polyline ROI of appropriate thickness. We used “Selection->Straighten” function of ImageJ on original z-stacks to generate B-spline interpolated stacks, so a dendrite became straight and oriented along X axis. From those stacks we generated a resliced stack in the plane perpendicular to dendrite’s axis (YZ) for the analysis of radial intensity distribution in the cross section (Fig.1A-B). To find the boundary outline of the dendrite in each slice, we used custom written set of ImageJ macros allowing semi-automated analysis [36]. The process consisted of two stages: finding the bounding rectangle encompassing the dendrite’s intensity and building a smooth closed spline (approximately in the shape of an oval, see below). Illustration of full analysis workflow is presented in Supplementary Movie 1.

First, we calculated a center of mass (based on intensity) coordinates *x*_c_, *y*_c_ for tyrosinated (STED data) or total tubulin (ExM) channels. Then we specified a rectangular ROI *R* of maximum area under conditions that it was still located inside the image and that its center was positioned at the center of mass. In the next step, we progressively downsized the rectangle from each side to find the position where edge’s intensity becomes equal to some threshold value (see below). We describe it here for the right side, but the same procedure was applied to all sides. Given an initial rectangle *R* of width *w*, height *h* and top left corner coordinates *x*_R_, *y*_R_, we built a set of rectangles with the width *w*_i_ ranging from *w*/2 to *w* (with a step size of one pixel) with the same height *h* and same and fixed position of the top left corner. For each rectangle from this set we measured the integrated intensity, providing the integrated intensity as a function of width *I*_int_(*w*_i_). The intensity of the edge *I*_e_(*w*_i_) was calculated as a derivative of this function, i.e. *I*_e_(*w*_i_)=*I*_int_(*w*_i_) – *I*_int_(*w*_i+1_) and normalized by its maximum and minimum value. A typical shape of *I*_e_(*w*_i_) represents a peak around *w*/2 (a center of dendrite/rectangle) that is gradually decaying towards periphery. For the first image in the resliced stack, by decreasing the value of *w*_i_ starting from *w*, we found the first value of *I*_e_(*w*_i_) that exceeds a threshold normalized intensity value of *I*_thr_ and its corresponding width *w*_RB_. The coordinate of the right boundary (RB) was calculated as *x*_RB_=*x*_R_ +*w*_RB_. The threshold intensity value *I*_thr_ was in the range of 0.2-0.4 and chosen for each first image in a stack manually to provide the values of *x*_RB_ corresponding to the visual boundary of dendrite’s intensity. The procedure was repeated for all other sides of the rectangle *R*, providing coordinates of left *x*_LB_, top *y*_TB_ and bottom *y*_BB_ boundaries. For horizontal boundaries, the width was kept the same and the position of the opposite edge was kept constant while building *I*_int_(*h*_i_). Using the newly found coordinates of the boundaries, we thereby built updated rectangle *R*_B_ encompassing dendrite’s cross section.

This method worked robustly in many cases, but it failed in the presence of axons that were often wrapped around a dendrite. In the YZ plane those axons produced additional fluorescent spots next to dendrite cross section that were included into rectangle *R*_B_. In the shape of *I*_e_(*w*_i_) curve they manifest themselves as additional local peaks. Therefore procedure of finding *R*_B_ from initial rectangle *R* for all other images in the resliced YZ stack (apart from the first) was modified. In these cases, we scanned *I*_e_(*w*_i_) by both decreasing and increasing values of *w*_i_ in the *w*/2 to *w* range. During a scan, we recorded all values of *w*_i_ that corresponded to each threshold *I*_thr_ from the set of 0.1 to 0.5 with a step of 0.1. From these we calculated a set of candidate right boundary positions, from which we chose the one that is closest to the corresponding boundary from the previous image in the stack. This value was recorded as the new edge of *R*_B_ rectangle at the current image. The procedure was repeated for each edge and after finding boundary rectangles for the whole stack, they were inspected and corrected manually.

To build a closed smooth spline contour around the irregular shaped dendrite’s cross section, in addition to vertical and horizontal boundaries, we also determined diagonal boundary points. For that we built an intensity profile along the 20-40 pixels wide line ROI connecting left top and right bottom corners of the rectangle *R*_B_. After normalization of intensity to minimum and maximum, we found coordinates of two points on the both halves of line where intensity is closest to 0.15-0.2 of its maximum value, denoted (*x*_LD1_, *y*_LD1_) and (*x*_LD2_, *y*_LD2_). The same procedure was performed on the diagonal segment connecting left bottom and top right corners of rectangle RB, providing points (*x*_RD1_, *y*_LD1_) and (*x*_RD2_, *y*_RD2_). The ordered set of eight points with coordinates (*x*_LB_, *y*_c_), (*x*_LD1_, *y*_LD1_), (*x*_c_, *y*_TB_), (*x*_RD1_, *y*_LD1_), (*x*_RB_, *y*_c_), (*x*_LD2_, *y*_LD2_), (*x*_c_, *y*_BB_), (*x*_RD2_, *y*_RD2_) was used to construct smooth closed spline boundary *C* passing through all of them (ImageJ functions *makePolygon* and “*Fit Spline*”). The final outlines for each image were inspected visually and if necessary, corrected manually.

In addition, for some ExM data profiles with low background we used an alternative, faster algorithm to find the boundary. We detected a set of points representing local fluorescence intensity maxima in each YZ slice (corresponding to MTs cross-sections). Using this set, we built a convex hull and constructed a spline from it. Again, we manually checked and corrected generated outlines.

To build the radial intensity distribution, for each image we iteratively reduced the contour *C* with steps of one pixel using ImageJ function “Enlarge ROI” with negative values (it uses Euclidean distance map threshold), while measuring its area *S*_k_ and integrated intensity *IC*_k_ (where index *k* denotes the step). We calculated the average intensity *MI*_k_ of each contour in the shrinking series as derivative:

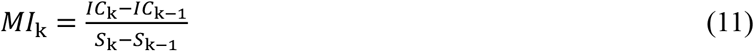

To get the radial distribution, for each *k* step we recalculated area *S*_k_ to radius using the formula 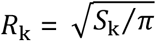 and normalized it by maximum value. Finally, to get a probability density function *ρ*(*R*), we normalized *MI*(*R*) by the area under the curve.

### Decomposition of radial intensities

For the decomposition of total tubulin radial density *ρ*_Tot_(*R*) as a weighted sum of tyrosinated *ρ*_Tyr_(*R*) and acetylated *ρ*_Ac_(*R*) densities (Fig.4G) we minimized the mean square error MSE(*w*_Tyr_,*w*_Ac_) between two curves:

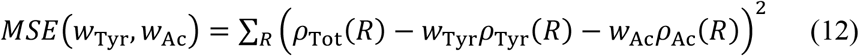

where *W*_Tyr_ and *W*_Ac_ correspond to the weights of tyrosinated and acetylated densities. By taking the derivatives of (12) and making them equal to zero, the solution can be found in a closed form:

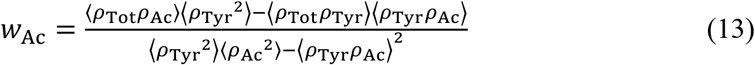

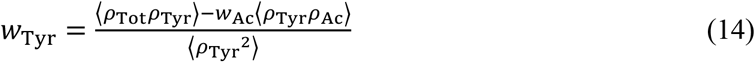

where angle brackets denote averaging over the whole radius range. It must be noted, that even without addition of a stronger assumption *W*_Tyr_ + *W*_Ac_ = 1, our analysis provided values that satisfy this relation.

### Single microtubule intensity estimate in FlipExM YZ stacks

The cross sections of microtubules in YZ FlipExM appeared as a set of fluorescent spots (Fig.5B). For intensity analysis we used ComDet v.0.5.3 plugin for ImageJ (https://github.com/ekatrukha/ComDet archived on Zenodo repository https://doi.org/10.5281/zenodo.4281064), which reports spot area, width, height in the detected channel and quantifies the background corrected integrated intensity in all three channels. To filter out possible microtubule bundles, we applied a lower bound threshold of 0.8 on the spot “roundness” *θ*, expressed as:

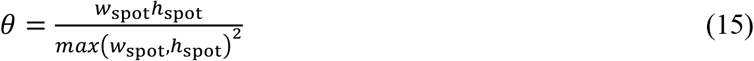

where *w*_spot_ and *h*_spot_ correspond to spot width and height. For spot areas detected per channel and per dendrite, we performed an MLE fit to the normal distribution and obtained estimates for the mean *S*_mean_ and standard deviation *σ*_area_, which were used to filter out spots with areas outside the inclusion range with lower bound max(*Q*_1_, *S*_mean_ – *σ*_area_) and upper bound (*S*_mean_+*σ*_area_), see Fig.5C,D. For the lower bound, *Q*_1_ stands for 25^th^ percentile, it was added to robustly remove false positives.

After the “roundness” and area filters, an estimation of average single microtubule intensity was performed in a similar way as in Fig.2, i.e. by fitting a sum of two Gaussians (Eq.(3)) to the histogram of intensity distributions (Fig.5E, S4A,B). For each slice of YZ stack we calculated normalized integrated intensity in each channel as a sum of all spot’s intensities divided by a single microtubule intensity derived from the fit. These intensity values were averaged over all slices for each channel and dendrite and used to calculate microtubule numbers according to Eq.(6)-(8) (as in Fig.1). The average level of tyrosination of microtubules detected in the acetylated channel *α* and the average level of acetylation of microtubules detected in the tyrosinated channel *β* was calculated per dendrite by pulling all filtered spot detections together (Fig.5F,G)

## SUPPLEMENTAL FIGURES

**Figure S1.**
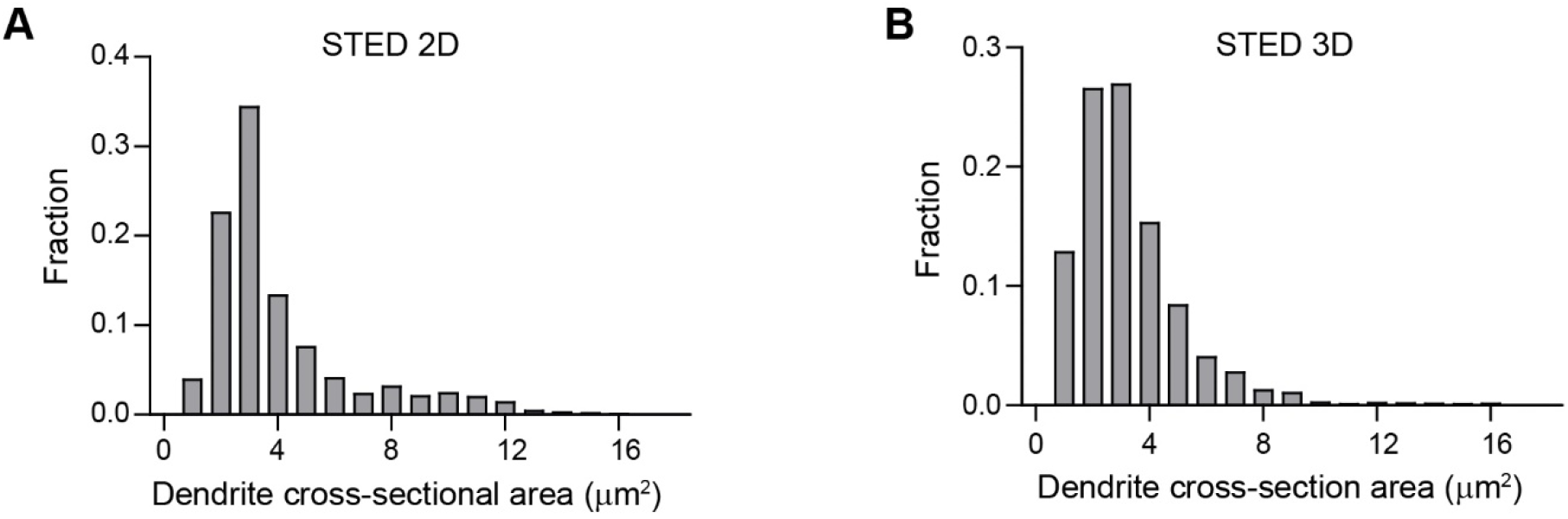
(A)-(B) Distribution of dendrite’s cross section area for the data shown in Fig.1(F)-(G).

**Figure S2.**
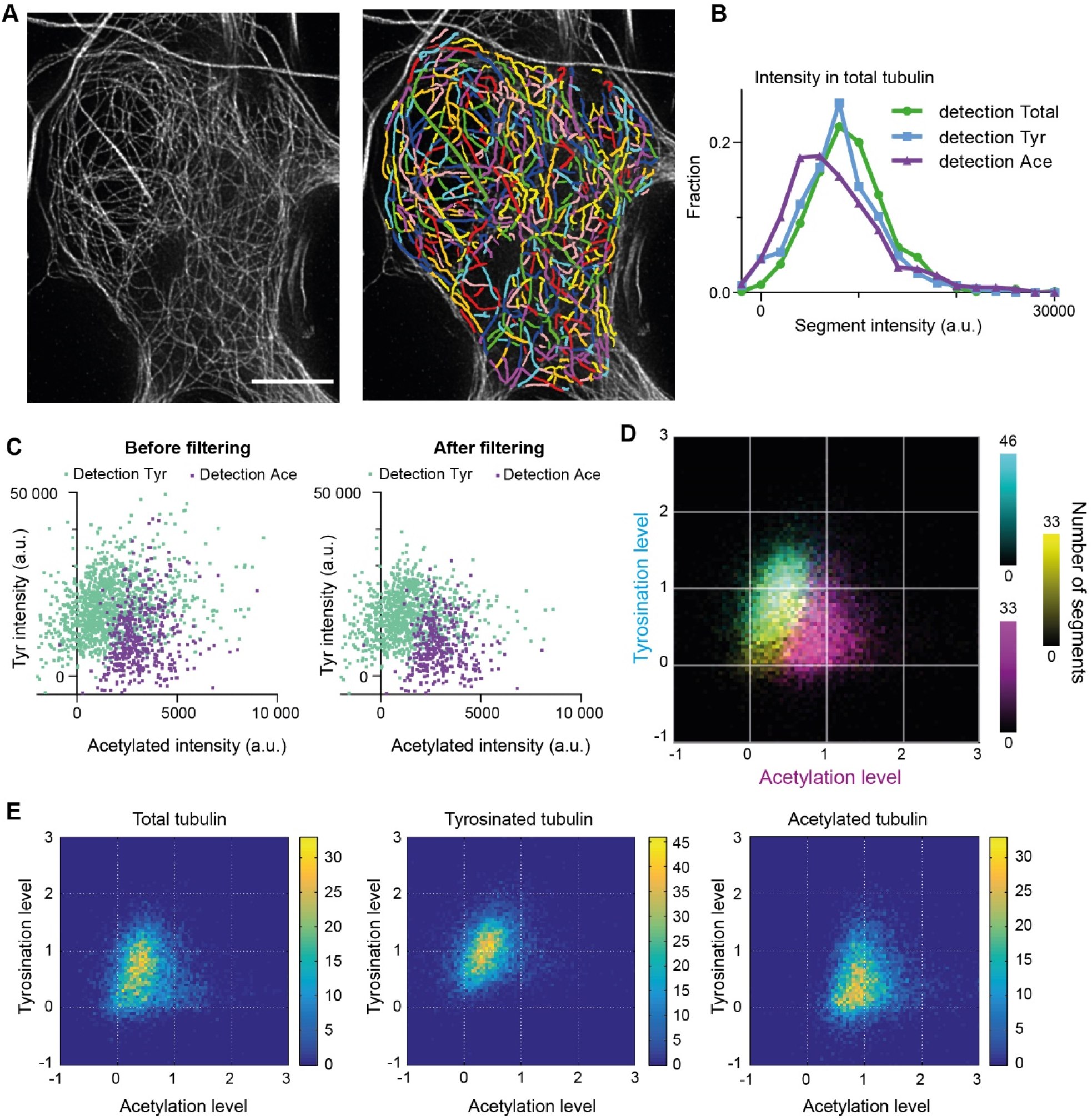
(A) Zoom-in of cell body area in total tubulin channel from Fig.2B, showing individual microtubule segments detections in different colors. (B) Histograms of background corrected integrated intensities of individual microtubule segments from the cell shown in Fig.2B, quantified in total tubulin channel and detected in total (green, n=660), tyrosinated (cyan, n=630) and acetylated (magenta, n=446) channels. (C) Background corrected integrated intensities of individual microtubule segments detected in tyrosinated (cyan) or acetylated (magenta) channels before (left) and after (right) threshold filtering (Fig.2E) in total tubulin channel (data for the cell shown in Fig.2B). (D) Three color heatmap overlay of normalized intensity distributions of single microtubule segments detected in total tubulin, (yellow, n=9050), tyrosinated (cyan, n=10281 segments) and acetylated (magenta, n=9369 segments) channels. (22 cells, N=2 independent experiments). (E) Heatmaps of normalized intensities in tyrosinated and acetylated channels for the detections performed in the specified channel built separately (data is the same as in (D)).

**Figure S3.**
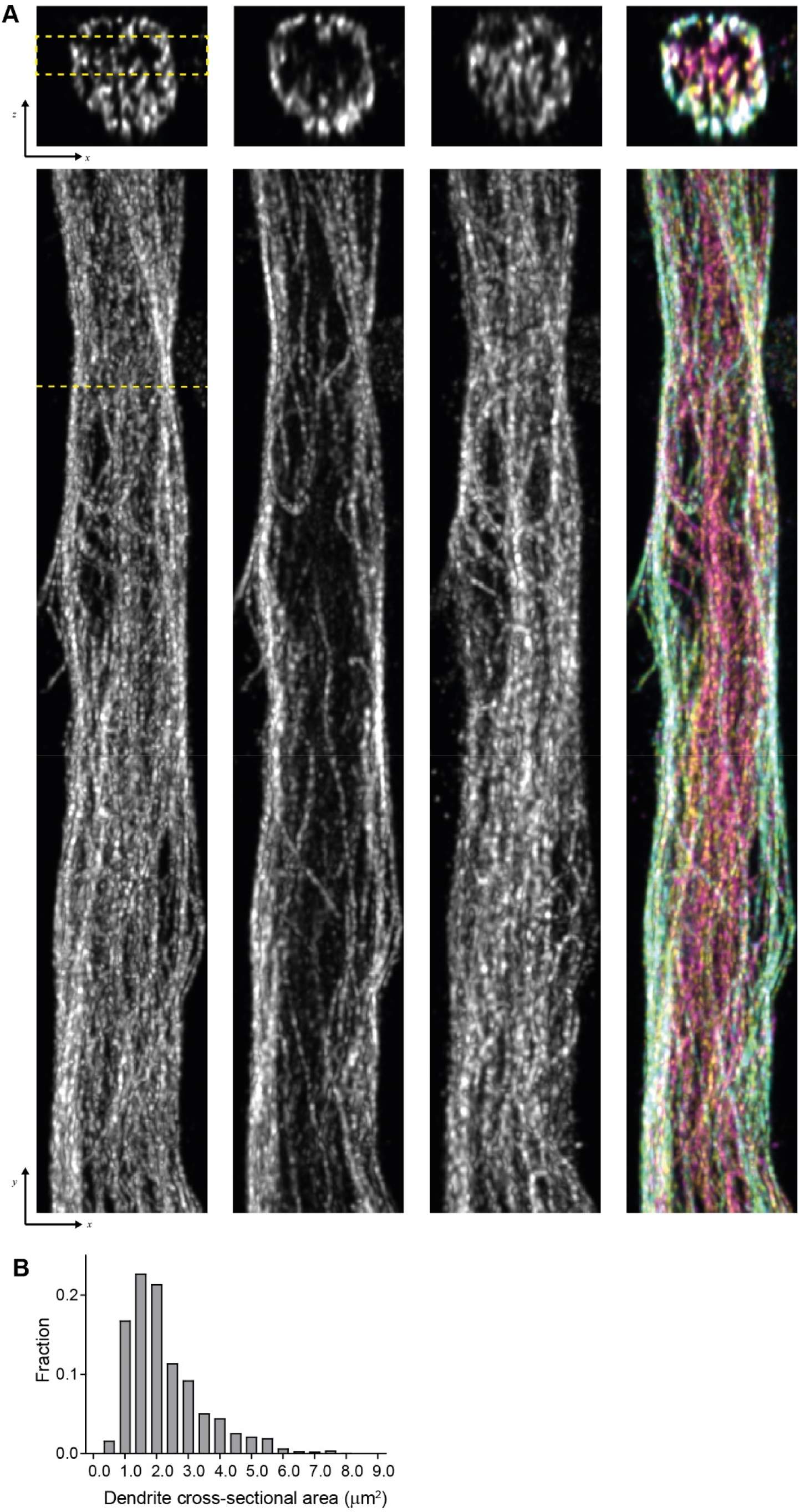
(A) (top) Representative single YZ cross section of the dendrite from Fig. 4A and (bottom) corresponding maximum intensity XY projections of subvolume marked by dashed yellow lines. Scale bar 0.5 μm (physical size post-expansion 2.07 μm). (B) Distribution of dendrite’s cross section area for the data shown in Fig.4(E)-(F).

**Figure S4.**
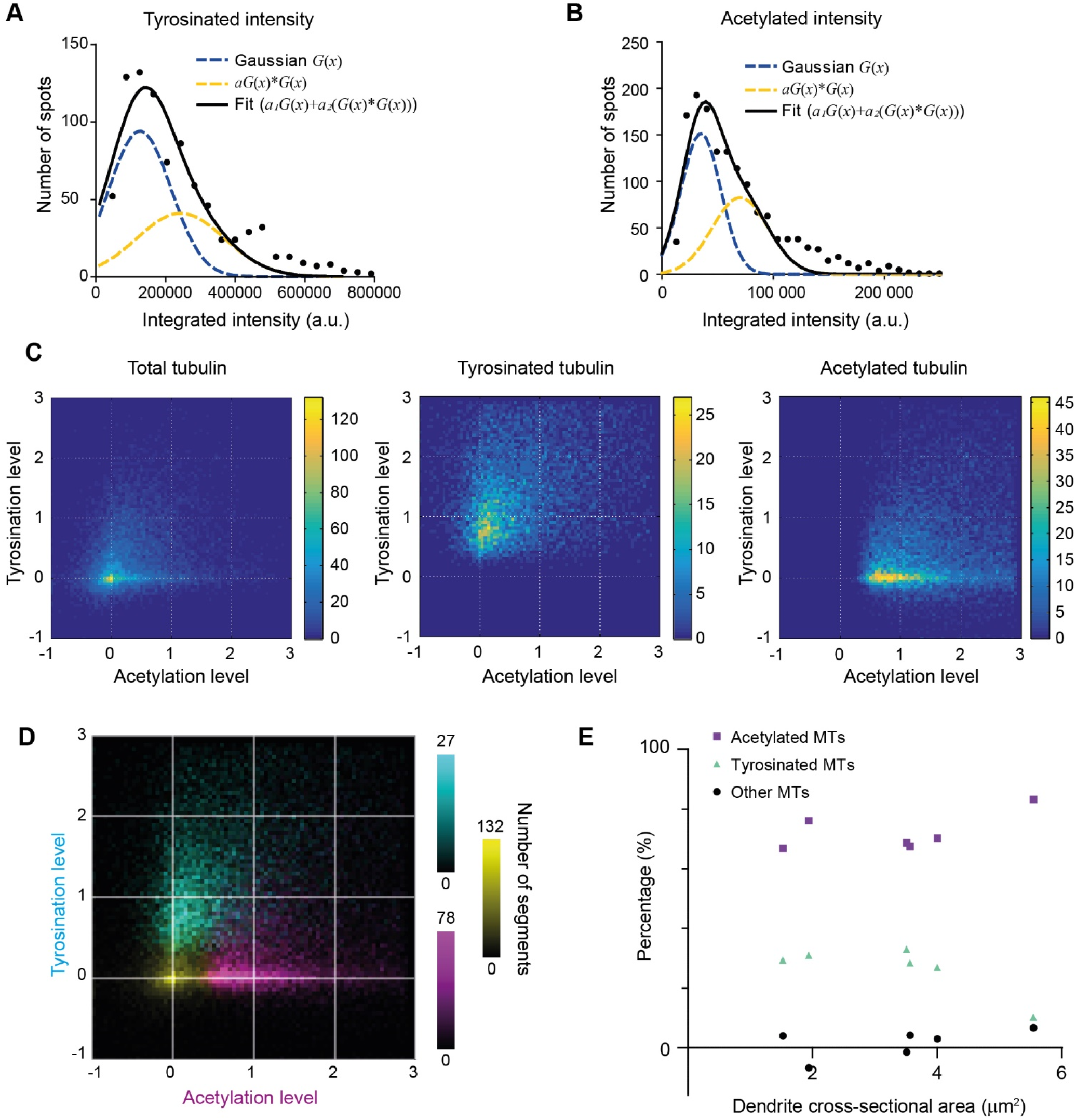
(A)-(B) Histograms of background subtracted integrated intensity of individual microtubules cross sections detected and quantified in tyrosinated (A) or acetylated (B) tubulin channel for the dendrite shown in (B) (black dots, n=870 and n=1441). The solid black line shows the fit of sum of two Gaussian functions: the first corresponds to a single microtubule cross section intensity distribution (dashed green line) and the second Gaussian corresponds to the double cross section intensity distribution, i.e. first Gaussian convoluted with itself (dashed blue line). (C) Heatmaps of normalized single microtubule cross section intensities in tyrosinated and acetylated channels for the detections performed in the specified channel built separately (data is the same as in (D)). (D) Three color heatmap overlay of normalized intensity distributions of single microtubule cross sections detected in total (yellow, n=20398), tyrosinated (cyan, n=8642) and acetylated (magenta, n=12552) tubulin channels (6 cells, N=2 independent experiments). Percentage of tyrosinated, acetylated and non-modified microtubules as a fraction of total microtubule number per dendrite as a function of dendrite’s cross section area (6 cells, N=2 independent experiments).

## SUPPLEMENTARY MOVIES

**Supplementary Movie 1**. Illustration of radial intensity distribution analysis in dendrites acquired using STED microscopy.

**Supplementary Movie 2**. 3D volumetric rendering of a dendrite imaged using ExM (same as in Fig.4A). Scale bar corresponds to the physical post expansion size.

**Supplementary Movie 3**. Illustration of gel sample reorientation and microscope’s PSF for FlipExM imaging.

**Supplementary Movie 4**. 3D volumetric rendering of a dendrite imaged using FlipExM (same as in Fig.5B). Scale bar corresponds to the physical post expansion size.

## REFERENCES

1. Stiess, M. and F. Bradke, Neuronal polarization: the cytoskeleton leads the way. Developmental neurobiology, 2011. 71(6): p. 430–444.

2. Bentley, M. and G. Banker, The cellular mechanisms that maintain neuronal polarity. Nature reviews. Neuroscience, 2016. 17(10): p. 611–622.

3. Kapitein, Lukas C. and Casper C. Hoogenraad, Building the Neuronal Microtubule Cytoskeleton. Neuron, 2015. 87(3): p. 492–506.

4. Burute, M. and L.C. Kapitein, Cellular Logistics: Unraveling the Interplay Between Microtubule Organization and Intracellular Transport. Annual Review of Cell and Developmental Biology, 2019. 35(1): p. 29–54.

5. Hirokawa, N., S. Niwa, and Y. Tanaka, Molecular Motors in Neurons: Transport Mechanisms and Roles in Brain Function, Development, and Disease. Neuron, 2010. 68(4): p. 610–638.

6. Kapitein, L.C. and C.C. Hoogenraad, Which way to go? Cytoskeletal organization and polarized transport in neurons. Molecular and Cellular Neuroscience, 2011. 46(1): p. 9–20.

7. Tempes, A., et al., Role of dynein–dynactin complex, kinesins, motor adaptors, and their phosphorylation in dendritogenesis. Journal of Neurochemistry, 2020. 155(1): p. 10–28.

8. Atherton, J., A. Houdusse, and C. Moores, MAPping out distribution routes for kinesin couriers. Biology of the Cell, 2013. 105(10): p. 465–487.

9. Sirajuddin, M., L.M. Rice, and R.D. Vale, Regulation of microtubule motors by tubulin isotypes and post-translational modifications. Nature cell biology, 2014. 16(4): p. 335–344.

10. Jurriens, D., et al., Mapping the neuronal cytoskeleton using expansion microscopy, in Methods in Cell Biology. 2020, Academic Press.

11. Monroy, B.Y., et al., A Combinatorial MAP Code Dictates Polarized Microtubule Transport. Developmental Cell, 2020. 53(1): p. 60-72.e4.

12. Park, J.H. and A. Roll-Mecak, The tubulin code in neuronal polarity. Current Opinion in Neurobiology, 2018. 51: p. 95–102.

13. Janke, C. and M.M. Magiera, The tubulin code and its role in controlling microtubule properties and functions. Nature Reviews Molecular Cell Biology, 2020. 21(6): p. 307–326.

14. Cai, D., et al., Single Molecule Imaging Reveals Differences in Microtubule Track Selection Between Kinesin Motors. PLOS Biology, 2009. 7(10): p. e1000216.

15. Dunn, S., et al., Differential trafficking of Kif5c on tyrosinated and detyrosinated microtubules in live cells. Journal of Cell Science, 2008. 121(7): p. 1085.

16. Guardia, Carlos M., et al., BORC Functions Upstream of Kinesins 1 and 3 to Coordinate Regional Movement of Lysosomes along Different Microtubule Tracks. Cell Reports, 2016. 17(8): p. 1950–1961.

17. Tas, R.P., et al., Differentiation between Oppositely Oriented Microtubules Controls Polarized Neuronal Transport. Neuron, 2017. 96(6): p. 1264-1271.e5.

18. Lipka, J., et al., Microtubule-binding protein doublecortin-like kinase 1 (DCLK1) guides kinesin-3-mediated cargo transport to dendrites. The EMBO Journal, 2016. 35(3): p. 302–318.

19. Nekooki-Machida, Y. and H. Hagiwara, Role of tubulin acetylation in cellular functions and diseases. Medical molecular morphology, 2020. 53(4): p. 191–197.

20. Roll-Mecak, A., How cells exploit tubulin diversity to build functional cellular microtubule mosaics. Current Opinion in Cell Biology, 2019. 56: p. 102–108.

21. Baas, P.W., et al., Polarity orientation of microtubules in hippocampal neurons: uniformity in the axon and nonuniformity in the dendrite. Proceedings of the National Academy of Sciences, 1988. 85(21): p. 8335.

22. Kubota, Y., et al., Conserved properties of dendritic trees in four cortical interneuron subtypes. Scientific Reports, 2011. 1(1): p. 89.

23. Steger, C., An unbiased detector of curvilinear structures. IEEE Transactions on Pattern Analysis and Machine Intelligence, 1998. 20(2): p. 113–125.

24. Zimmermann, T., Spectral Imaging and Linear Unmixing in Light Microscopy, in Microscopy Techniques: -/-, J. Rietdorf, Editor. 2005, Springer Berlin Heidelberg: Berlin, Heidelberg. p. 245–265.

25. Tillberg, P.W., et al., Protein-retention expansion microscopy of cells and tissues labeled using standard fluorescent proteins and antibodies. Nature Biotechnology, 2016. 34(9): p. 987–992.

26. Gao, M., et al., Expansion Stimulated Emission Depletion Microscopy (ExSTED). ACS Nano, 2018. 12(5): p. 4178–4185.

27. Schulze, E. and M. Kirschner, Dynamic and stable populations of microtubules in cells. The Journal of cell biology, 1987. 104(2): p. 277–288.

28. van de Willige, D., C.C. Hoogenraad, and A. Akhmanova, Microtubule plus-end tracking proteins in neuronal development. Cellular and Molecular Life Sciences, 2016. 73(10): p. 2053–2077.

29. Akhmanova, A. and M.O. Steinmetz, Control of microtubule organization and dynamics: two ends in the limelight. Nature Reviews Molecular Cell Biology, 2015. 16(12): p. 711–726.

30. Esteves da Silva, M., et al., Positioning of AMPA Receptor-Containing Endosomes Regulates Synapse Architecture. Cell Reports, 2015. 13(5): p. 933–943.

31. Jaworski, J., et al., Dynamic Microtubules Regulate Dendritic Spine Morphology and Synaptic Plasticity. Neuron, 2009. 61(1): p. 85–100.

32. McVicker, D.P., et al., Transport of a kinesin-cargo pair along microtubules into dendritic spines undergoing synaptic plasticity. Nature Communications, 2016. 7(1): p. 12741.

33. Hu, X., et al., Activity-Dependent Dynamic Microtubule Invasion of Dendritic Spines. The Journal of Neuroscience, 2008. 28(49): p. 13094.

34. Schätzle, P., et al., Activity-Dependent Actin Remodeling at the Base of Dendritic Spines Promotes Microtubule Entry. Current Biology, 2018. 28(13): p. 2081-2093.e6.

35. Kapitein, L.C., K.W. Yau, and C.C. Hoogenraad, Microtubule Dynamics in Dendritic Spines. Microtubules: In Vivo, 2010. 97: p. 111–132.

36. Katrukha, E.A., et al., Quantitative mapping of dense microtubule arrays in mammalian neurons. Datasets, code and figures. Available at https://doi.org/10.6084/m9.figshare.c.5306546.v1. 2021.

